# It’s about time: Analysing an alternative approach for reductionist modelling of linear pathways in systems biology

**DOI:** 10.1101/781708

**Authors:** Niklas Korsbo, Henrik Jönsson

**Affiliations:** The Sainsbury Laboratory, University of Cambridge, Cambridge, United Kingdom; Department of Applied Mathematics and Theoretical Physics, University of Cambridge, Cambridge, United Kingdom; Department of Astronomy and Theoretical Physics, Computational Biology and Biological Physics, Lund University, Lund, Sweden

## Abstract

Thoughtful use of simplifying assumptions is crucial to make systems biology models tractable while still representative of the underlying biology. A useful simplification can elucidate the core dynamics of a system. A poorly chosen assumption can, however, either render a model too complicated for making conclusions or it can prevent an otherwise accurate model from describing experimentally observed dynamics.

Here, we perform a computational investigation of linear pathway models that contain fewer pathway steps than the system they are designed to emulate. We demonstrate when such models will fail to reproduce data and how detrimental truncation of a linear pathway leads to detectable signatures in model dynamics and its optimised parameters.

An alternative assumption is suggested for simplifying linear pathways. Rather than assuming a truncated number of pathway steps, we propose to use the assumption that the rates of information propagation along the pathway is homogeneous and instead letting the length of the pathway be a free parameter. This results in a three-parameter representation of arbitrary linear pathways which consistently outperforms its truncated rival and a delay differential equation alternative in recapitulating observed dynamics.

Our results provide a foundation for well-informed decision making during model simplifications.

**Author summary:** Mathematical modelling can be a highly effective way of condensing our understanding of biological processes and highlight the most important aspects of them. Effective models are based on simplifying assumptions that reduce complexity while still retaining the core dynamics of the original problem. Finding such assumptions is, however, not trivial.

In this paper, we explore ways in which one can simplify long chains of simple reactions wherein each step is linearly dependent on its predecessor. After generating synthetic data from models that describe such chains in explicit detail, we compare how well different simplifications retain the original dynamics. We show that the most common such simplification, which is to ignore parts of the chain, often renders models unable to account for time delays. However, we also show that when such a simplification has had a detrimental effect, it leaves a detectable signature in its optimal parameter values. We also propose an alternative assumption which leads to a highly effective model with only three parameters. By comparing the effects of these simplifying assumptions in thousands of different cases and for different conditions we are able to clearly show when and why one is preferred over the other.

## Introduction

Biochemical reaction networks are often complicated and any attempt to describe them using mathematical models relies heavily on simplifying assumptions (1). Effective models are often built upon simplifying assumptions that avoids over-fitting by using as few free parameters as possible while still capturing the main properties of the biological system (2, 3). Thoughtful assumptions, as well as robust methods to identify parameter values, and (semi) global analysis of dynamical behaviour within model spaces, are all essential when evaluating models (4–7). However, assumptions that are beneficial in one setting may be detrimental in another and it is important, although non-trivial, to identify when this happens (8).

Multi-step processes are ubiquitous in biology. Examples are transcription and translation, where an RNA polymerase or a ribosome can perform thousands of sequential reactions before a protein is produced. Yet, in gene-regulatory networks, this is often reduced to a one or two-step reaction of a transcription factor which may lead to an mRNA before it leads to a finished protein (e.g. 1, 9–11). Another example is kinase cascades, where the product of a kinase triggers the action of downstream kinases (12). A well known such cascade can be found in the MAP kinases, which are triggered by the MAP kinase kinases, which in turn is triggered by the MAP kinase kinase kinases (13–18). There are also molecules which must undergo sequential multi-site phosphorylations before they are activated and can pass on any signalling (19, 20). This is, for example, important for the Drosophila circadian clock protein CLOCK who’s inactivity, activity, and degradation are thought to be governed by its sequential states of phosphorylation (21). Sequential multi-step reactions can also be important for signal perception and transduction pathways. One example is the receptor-like kinase FLAGELLIN SENSING 2 which, upon detecting of a pathogen, triggers a long chain of phosphotransfers, phosphorylations, and subcellular re-localisations that eventually leads to an immune response in *Arabidopsis thaliana* (22, 23). Another well studied system is the TGF*β* growth factor, which similarly triggers a sequence of phosphorylation steps before affecting the expression of downstream genes (24–26).

Linear pathways represent a large class of multi-step reactions which are both biologically relevant and theoretically approachable. Multi-step pathways can be represented as a chain of state changes where the activation rate of one state is dependent on the activity of the previous state. The mechanisms by which one active state regulates the activation of the next may be complicated but it can be useful to approximate these as being linear. This is partly since it is a minimally complex assumption and partly because many biochemical reactions appear to be linear as long as they operate in a weakly activated manner, far from saturation (12, 27).

Linear pathways are dynamically important and can entirely change the qualitative behaviour of a model. Their main effects are to supply signal amplification/dampening and to provide delays in the signalling (12, 28). The amplitude modulation of the signalling is governed by the ratio between the activation and inactivation rates; a pathway step will provide amplification if its activation rate exceeds its inactivation rate. The time-delay, on the other hand, is governed by the inactivation rates and by the length of the pathways (12, 27). These time-delays can have a significant impact on how a biological system works. A striking example of this is that delays are required for oscillations to be possible (29, 30).

The modelling of linear pathways pose a specific set of challenges. Full enumeration of the linear pathway greatly increases model complexity yet add disproportionately little in terms of dynamical range. However, even if the individual steps are of little dynamical significance, the aggregate effect of the full pathway may not be. There is therefore a need for a simplifying assumption which reduces the complexity of the linear pathway while still representing its total effect.

A common way of simplifying linear pathways is to ignore most of the reaction steps and assume that a model can recapitulate their effect using only one or a few steps (3). While this assumption is often implicit, it is easy to find examples where it has been used to simplify multi-step reactions such as protein production (e.g. 7, 31–33); protein-to-protein signalling networks (e.g. 7, 32–34); protein modifications such as phosphorylations, methylations, and ubiquitinations (e.g. 33, 34); and more. However, despite the frequent use of such pathway truncation, little effort has been made to understand its general consequences.

An alternative simplification is to represent the effect of the linear pathway using a fixed time-delay in the model. Focusing on this aspect of the linear pathway and assuming that all other aspects are negligible allows for a terse model description using delayed differential equations (DDEs) (35–38). However, it is not clear how such an assumption limits a model’s ability to recapitulate the dynamics of the full system.

A third simplification is to make use of a gamma distributed delay. This approach models the output of the linear pathway as the convolution between the input to the first pathway step and the probability density function of the gamma distribution. This convolution has been used to describe delays in a diverse set of processes, including: drug uptake (39, 40), circadian clocks (41–43), population dynamics (44) and even traffic jams (45). Similar to the fixed-delay approach, it is commonly used as a method to introduce a delay without explicit regard to what the underlying cause of that delay is. It can be derived from a chain of identical linear processes which indicates that it may be particularly relevant for linear pathways. However, an understanding of how well this simplification can represent a general linear pathway is still missing.

Here, we analyse the dynamical effect of linear pathways. First, we focus on what dynamical properties a model will be unable to reproduce when it is simplified using pathway truncation. This analysis led us to a diagnostic tool for revealing when such a model assumption has had detrimental effects.

Thereafter, we suggest the use of an alternative simplifying assumption and demonstrate its effectiveness. Rather than assuming a fixed (and truncated) pathway length, we assume a fixed rate of information propagation along a pathway of dynamic length. This leads to a three-parameter model which can recapture the dynamics of arbitrary linear pathways with high fidelity. The assumption allows for a direct derivation of the gamma distributed delay and it allows the model parameters to be anchored to the underlying biology. Furthermore, it outperforms the use of the reduced step approximation as well as the fixed-delay approximation and it provides a building block for an operational model inference approach (46).

## Results

### Linear pathway truncation causes different degrees of error for different underlying distributions of reaction rates

We set out to explore the dynamical consequences of misrepresenting the number of pathway steps in a model of a linear pathway. A main aim was to understand whether and how a short (truncated) linear pathway model fails to reproduce the dynamics generated by a longer pathway.

To investigate this, we first defined a model wherein a sequence of *n* states, with concentrations *X*_1_, *X*_2_, …, *X*_*n*_, each activates its successor. A step, *i*, in such a pathway is linear if

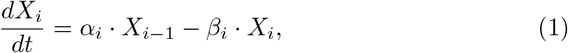

for some activation rate constant *α*_*i*_ and degradation/inactivation rate constant *β*_*i*_. We define the entire pathway as linear if this equation holds for all steps. In this study, we specifically focus on the relationship between an input and the output of a linear pathway and not on the relative concentrations along the different pathway steps. We can thus define a set of new parameters, such that scaling is done by a single parameter, *γ*, and the response rate of pathway step *i* to changes in the previous step is governed by a parameter *r*_*i*_ (Methods). This leads to a model that describes how an *n*-step linear pathway transforms an input signal, *I*(*t*), to an output, as defined by the concentration of the *n*-th step, which is given by

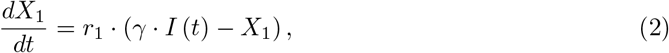

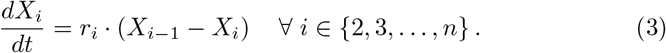

Synthetic data sets were generated using Eqs. 2-3 with different pathway lengths, *n*_*data*_ ∈ {1, 2, …, 50}. The response rates were drawn from a log-uniform distribution, 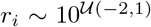, and the scaling parameters, *γ*, was set to one.

The effects of misrepresenting the pathway length in a model was tested on each set of synthetic data. Models with a fixed pathway step length (fixed-step models) of *n*_*model*_ = 1, 2, …, 5, respectively, were each treated with the same input and initial conditions that was used for data generation and had their parameters values (*γ*, *r*_*i*_ ) optimised to fit the output dynamics of that data (Methods). We first used the simplest possible model conditions to study how well a model could perform when *n*_*model*_ ≠ *n*_*data*_. Here, the pathway received no input (*I* (*t*) = 0) but was instead initialised with a non-zero concentration of the first, and only the first, pathway step. This could represent a sudden start of a reaction at *t* = 0 where *X*_1_ passes on its signalling while being exponentially depleted itself. In order to still have a parameter *γ* to scale the system and in order to ensure that the reaction rate parameters do not affect this scaling, we set the initial concentration to *X*_1_ (0) = *γr*_1_, which ensures that 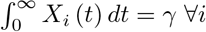.

We analysed how such models can reproduce the generated data and especially how the model/data fit depends on how many pathway steps were used to generate that data (Fig. 1 and Figs. S1-S4). When *n*_*model*_ ≥ *n*_*data*_, the model can perfectly reproduce the output dynamics, as expected, and provides a control for our numerical optimisation scheme (Fig. 1 and Figs. S1-S4). However, despite the perfection in the input-output correspondence, the fitted model and the data-generating model will not be precisely the same. While every optimal reaction rate in the fitted model will also be found in the data-generating linear pathway (Fig. 3b), they do not necessarily appear in the same order (Fig. 3c). The order of the response rates along the pathway does not matter for the output (27, 28). This means that optimising for an input-output relationship will not provide any means of correctly inferring which rate belongs to which step. Not only does the model correctly fit the data when *n*_*model*_ = *n*_*data*_, but also when *n*_*model*_ > *n*_*data*_ (Fig. 1 and Figs. S2-S4). During the optimisation procedure, ‘additional’ rates become fast enough to ‘instantaneously’ pass information from the previous step to the next (Fig. 3b), confirming that fast steps are less dynamically relevant than slow steps (1, 27).

**Figure 1:**
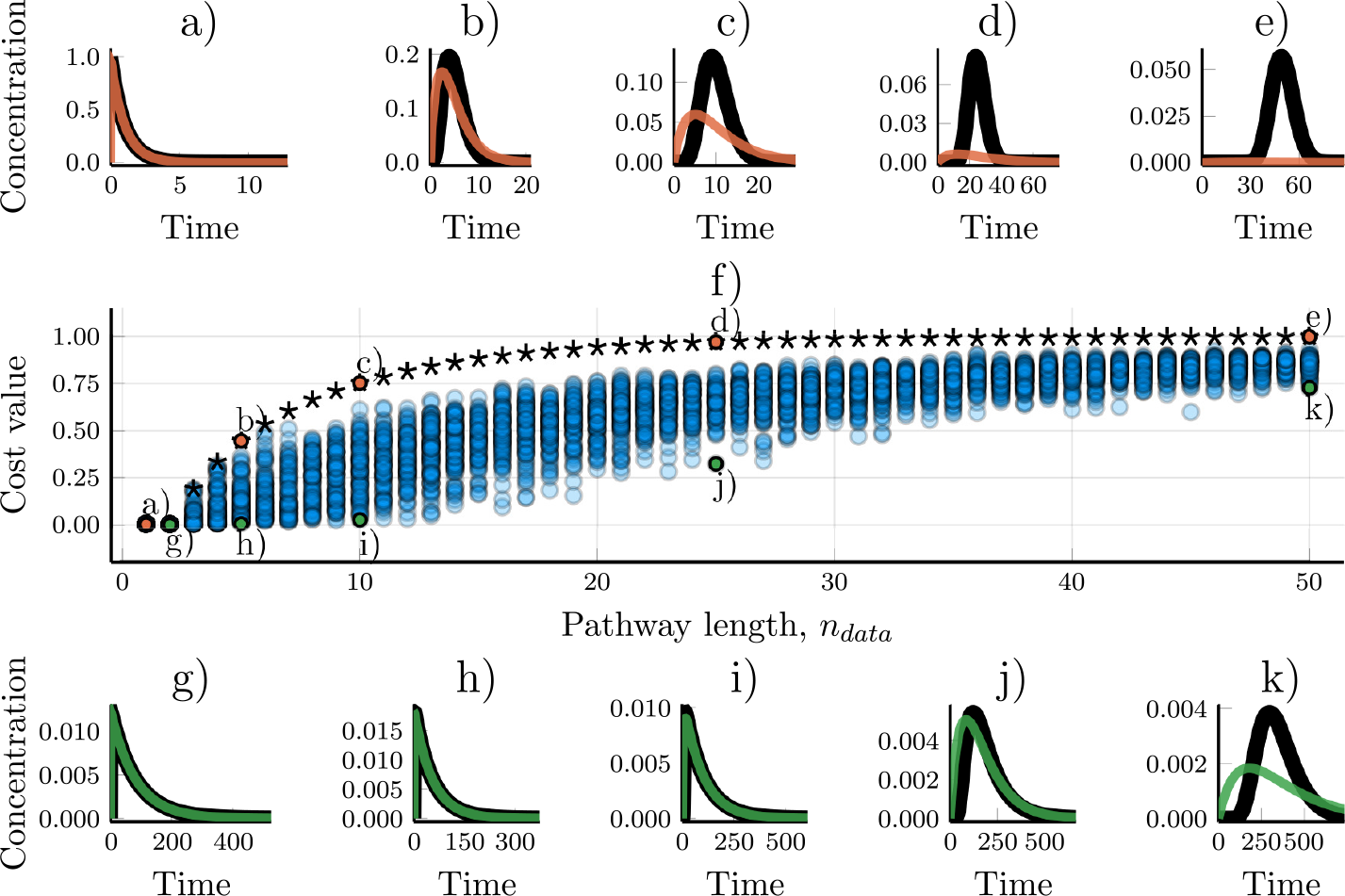
Modelling linear pathways using a truncated number of pathway steps. Two-step linear pathway models (Eqs. 2-3) were fitted towards synthetic data. The data shown was generated by networks of step-lengths varying from 1 to 50. a-e) The worst model/data fits for a given length, *n*_*data*_, of the model that generated the data. Black lines show the synthetic data while simulations of the fitted models are overlaid in colour. (f) The cost value for models optimised towards 5000 different sets of synthetic data. Stars are the cost values resulting from data wherein all the steps in the data-generating linear pathway have the same reaction rates, *r*_*i*_ = 1 ∀ *i*. The x-axis shows the number of steps in the models that were used to generate the data. g-k) Examples of the best model/data fits for different data pathway lengths, *n*_*data*_.

When the fitted model has fewer steps than the linear pathway which was used to generate the data, *n*_*model*_ < *n*_*data*_, the fit is no longer guaranteed to be perfect. Unsurprisingly, the model/data mismatch increases with the length of the pathway that generated the data and decreases with the length of the model used to fit the data (Fig. 1 and Figs. S1-S4).

There is a high variability in the ability of a truncated model to fit the output dynamics (Fig. 1 and Figs. S1-S4). While small models cannot in general represent arbitrary linear pathways, in some cases they are performing well. For example, a two-step model can be good at reproducing the dynamics of even a 10-step linear pathway (Fig. 1j), and a five-step model can accurately describe the dynamics of some 25-step pathways (Fig. S4k). The optimised model performance decreases with the homogeneity of the reaction rates of the data-generating pathway (Fig. 2). When all the response rates of the data-generating pathway are the same, the ability of models to fit the data quickly decreases with the length of the pathway (Fig. 2, cf. Figs. 1a-e, stars in Fig. 1f). Conversely, when the response rates are highly heterogeneous, even a heavily truncated model is able to fit data from a long pathway (Fig. 2), again indicating the different contributions of fast and slow steps to the resulting dynamics.

**Figure 2:**
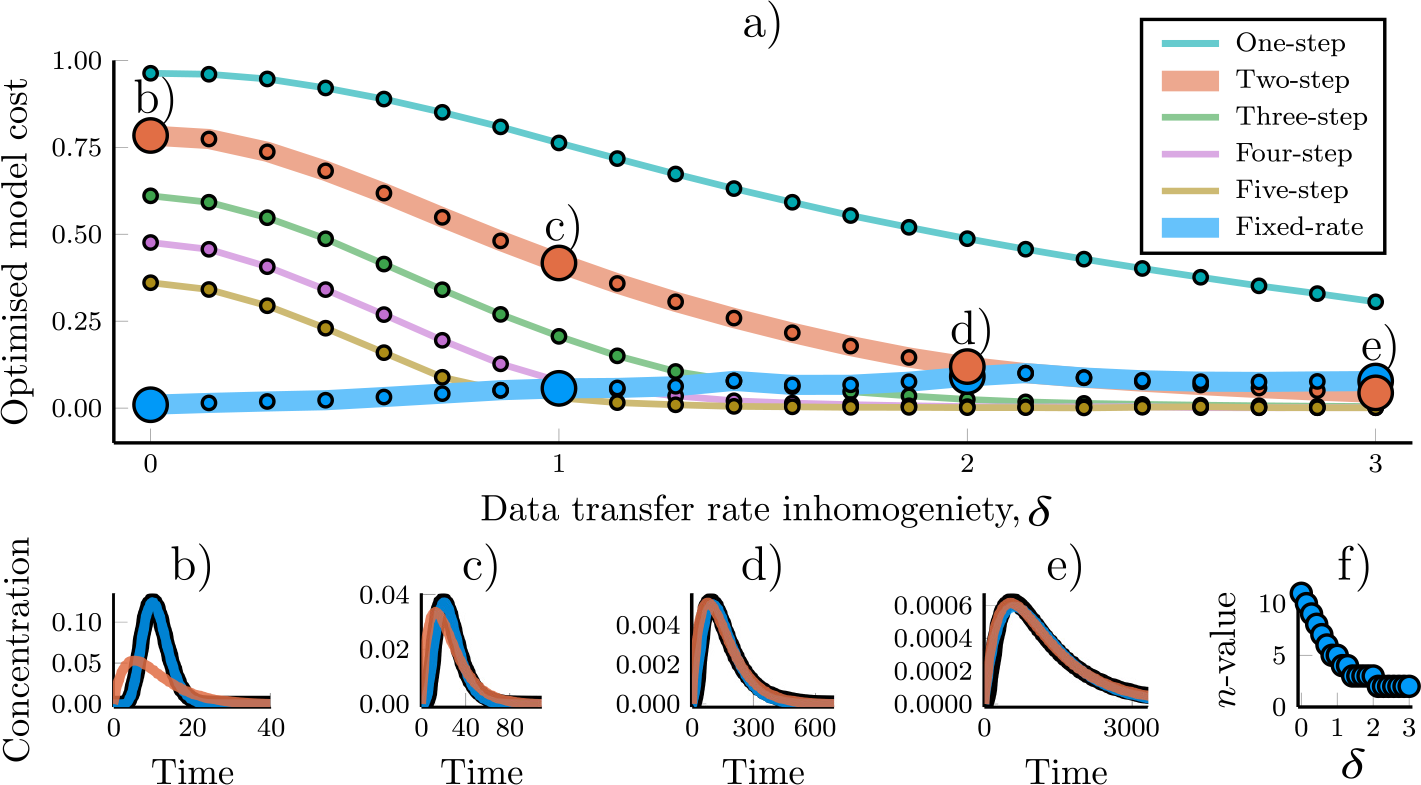
Response rate homogeneity of a linear pathway affect how well models can reproduce their dynamics. Models were fitted towards data that was generated with different levels of response rate inhomogeneity. The synthetic data was generated using an 11-step linear pathway model (Eqs. 2-3) with parameters *γ* = 1, 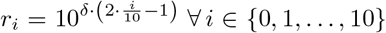 where *δ* is a parameter that governs the inhomogeneity of the rate parameters. For *δ* = 3, the rate parameters were thus logarithmically spread from 0.001 to 1000. a) The optimised cost values of different models, plotted against the inhomogeneity parameter, *δ*, used to generate the synthetic data. b-e) Samples of the model/data fit for the optimised parameter sets, as marked in a). The fixed-rate model (Eqs. 4-5) is compared with the two-step model (Eqs. 2-3) since they have the same number of free parameters. f) The optimised value for the fixed-rate model parameter *n*, depending on the inhomogeneity of the response rates.

**Figure 3:**
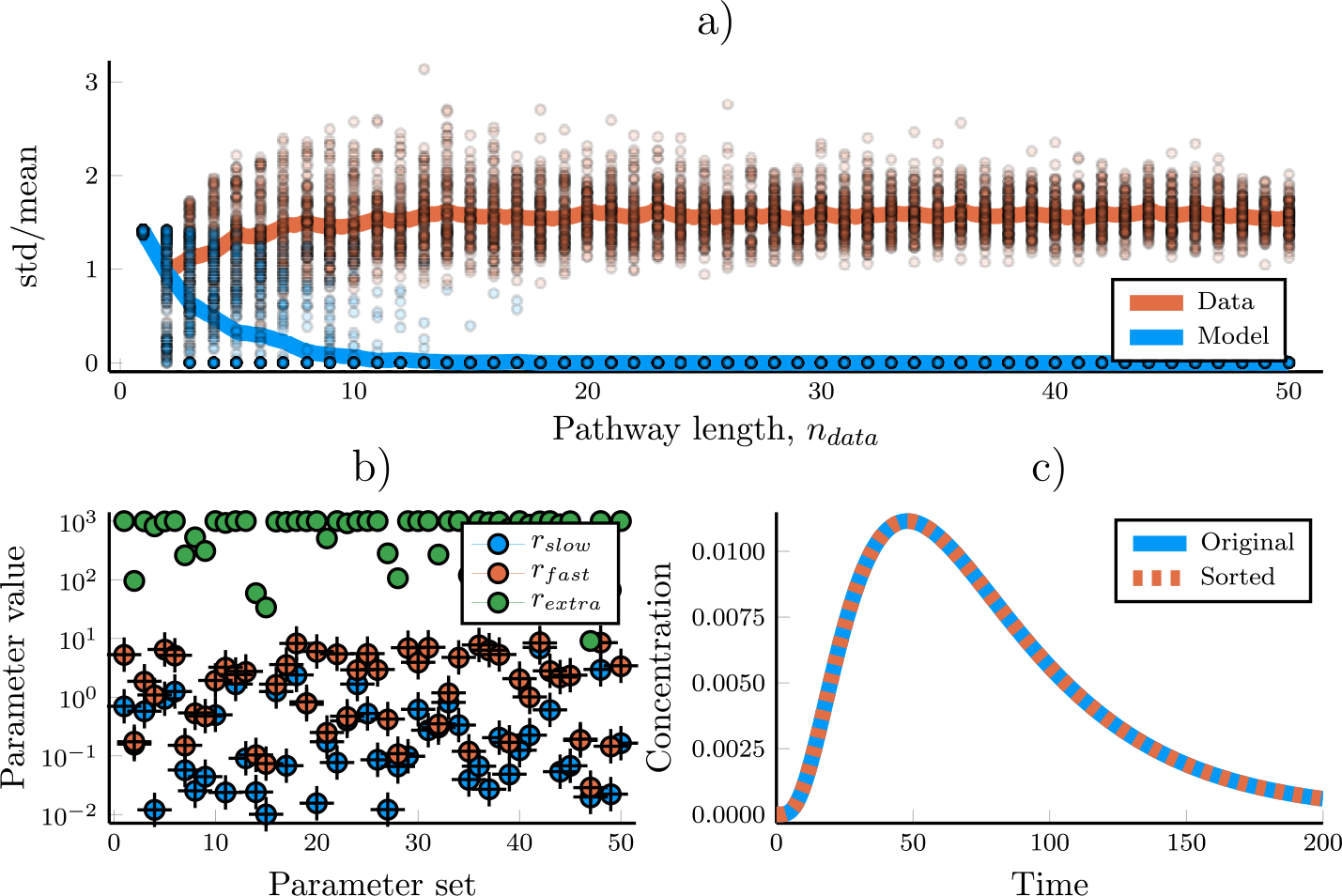
Analysis of optimised parameter values for the fixed-step models (Eqs. 2-3). a) The standard deviation, normalized by the mean, of the optimized two-step model parameters as well as for the parameters used to generate the synthetic data. Circles shows the standard deviation normalized by the mean for individual parameter sets and the solid lines are the mean of these values. b) The optimised parameters of a model with three steps, fitted against data generated using only two steps. The model parameter values are shown as circles, and the corresponding parameter values used to generate the data are shown as crosses. The green circles represent the model parameter values of the superfluous pathway step. c) The same model run twice, but with the order of its response rate parameters changed.

### Detrimentally truncated models of linear pathways can be identified by characteristic parameter values

While pathway length and response rate homogeneity are key determinants for whether the truncation of a linear pathway reduces a model’s accuracy, these features may often be unknown from experiments. Hence, it would be useful to find characteristics of the (simplified) model to quantitatively identify when it is performing badly.

One feature identified in our simulations is that the signalling peak in an over-simplified model has a lower amplitude and increased temporal width than that of the corresponding data (Fig. 1, Figs. S2-S4). This occurs when the model must compensate for the delay that a multi-step reaction causes. The compensation occurs in the form of slower response rates and that results in a less sharply defined output curve (cf. 12).

Possibly more useful, a detrimentally truncated model created a clear signature in the optimised parameter values in our test case. The standard deviation of the optimal response rate parameters decreases as *n*_*data*_ − *n*_*model*_ increases (Fig. 3a). When the model is much too small for recapitulating the data, all of its rate parameter values end up being the same (Fig. 1a). The variability in rate parameters hence provides a quantitative measure to detect detrimentally truncated linear pathways.

### An alternative assumption for model simplification improves predictability of pathway output dynamics

An alternative approach for parameter reduction is to assume that every path-way step has the same response rate (*r*_*i*_ = *r* ∀*i*), and to treat the number of pathway steps, *n*, as a free parameter. With this ‘fixed-rate’ assumption we can represent a linear pathway with the set of equations

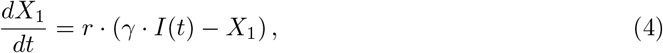

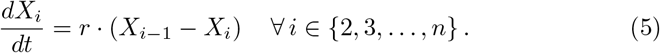

This simplified model has only three free parameters: *γ* for scaling, *n* for pathway length, and *r* for the response rates of the pathway steps.

While this assumption of homogeneous reaction rates is natural when the same process is repeated multiple times, such as a molecular motor walking along a microtuble, or for the assembly of monomers into a polymer, it is not true for most pathways. We analysed the effectiveness of the fixed-rate model by individually fitting its three parameters towards each of the synthetic data sets used above. The resulting model/data fits show that the fixed-rate assumption is indeed well suited for modelling linear pathways, even when the reaction rates of the data-generating network are highly heterogeneous (Fig. 4 and Fig. 2). Not only does the fixed-rate model outperform the two-step truncated model, which has the same number of parameters (Fig. 4), it also improves the prediction of pathway dynamics compared to the five-step (6 parameters) truncated model (cf. Fig. S4). Unlike the truncated, ‘fixed-step’, models, the fixed-rate model’s ability to fit data does not decrease with *n*_*data*_ (Fig. 4f). While the two-step model had costs (for definition, see Methods) ranging from 0 to almost 1, the fixed-rate model’s cost never exceeded 0.2 for our synthetic data sets. Indeed, the fixed-rate model proved more able to fit the data than the fixed-step models in a clear majority of cases (Figs. 4o and S6). This holds true for a wide range of different inputs to the linear pathway (Fig. 5). Strikingly, the fixed-rate model was almost unseperable from the output of the original model in most cases (blue and black lines in Fig. 5). The fixed-rate assumption is thus highly effective at simplifying a linear pathway, and the performance only decrease slightly when the underlying pathway has heterogeneous reaction rates (Fig. 2).

**Figure 4:**
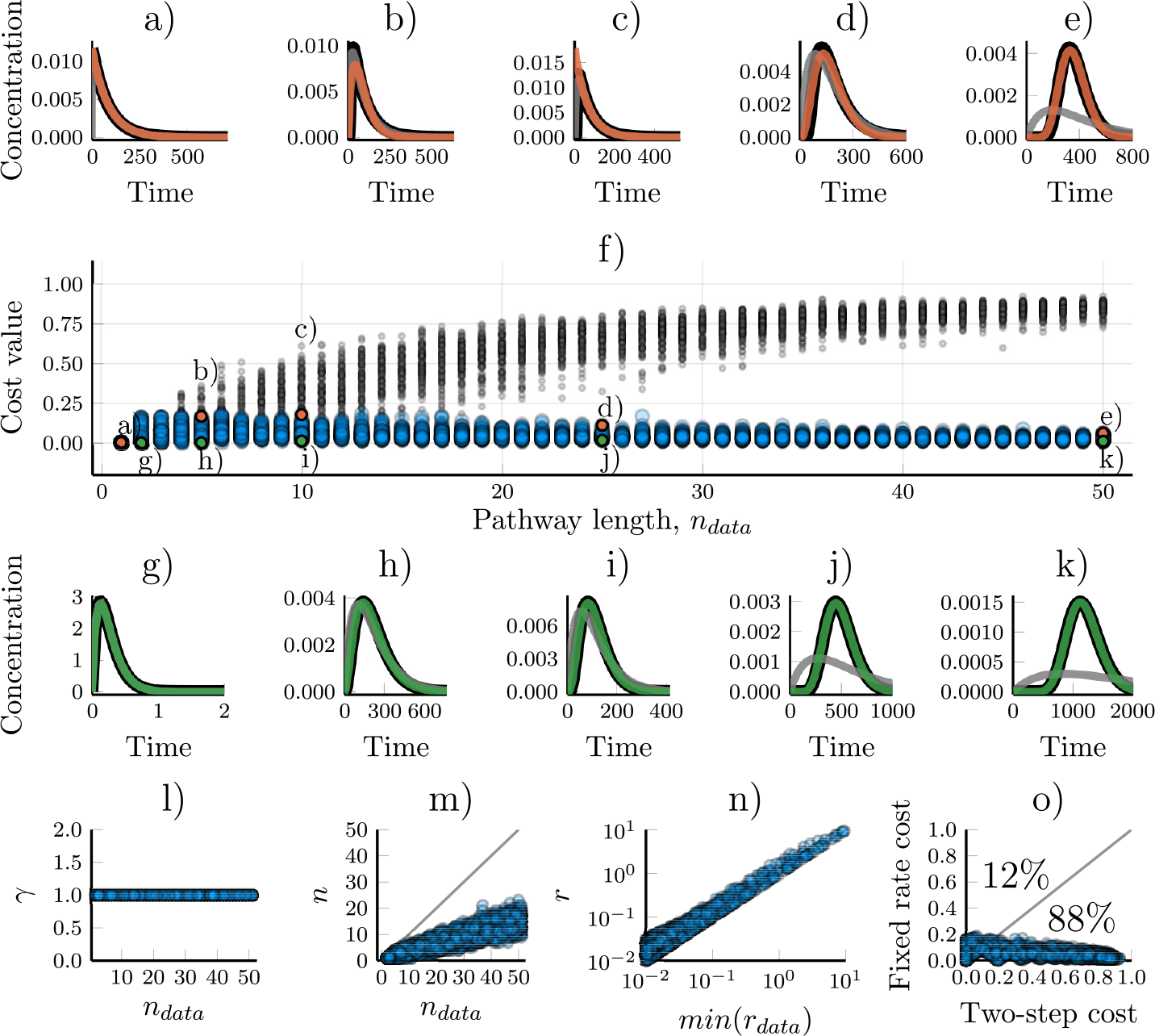
Modelling linear pathways using the fixed-rate assumption for simplification. The fixed-rate model (Eq. 4-5) was fitted towards synthetic data generated by networks of step-lengths varying from 1 to 50. The data sets are the same as those used for Figure 1. a-e) Examples of the worst model/data fits for the fixed-rate model. Orange lines show simulations of the fitted model and the black line shows the synthetic data. The grey line is the corresponding fit using the two-step truncated model on the same data set. f) The cost value for models optimised towards 5000 different sets of synthetic data. The x-axis shows the number of steps in the model which were used to generate the data. Blue circles are cost values for the fixed-rate model while grey dots are the cost values for a two-step truncated model. The parameter sets used in figures a-e and g-k are marked accordingly. g-k) Examples of the best model/data fits. The fixed-rate model (green lines) almost completely matches the data (black lines). l) The scaling parameter, *γ*, is accurately identified as 1 in all optimisations. m) The optimised values of *n* for different number of steps in the pathway underlying the data. n) A comparison between the optimised value of *r* and the smallest rate constant of the model that generated the data. o) A comparison of the cost values when using either the fixed-rate model or a two-step truncated model to fit the same data. Each circle represents a single synthetic data set. Percentages indicate how many of the data sets had a higher (worse) cost value for the respective models.

**Figure 5:**
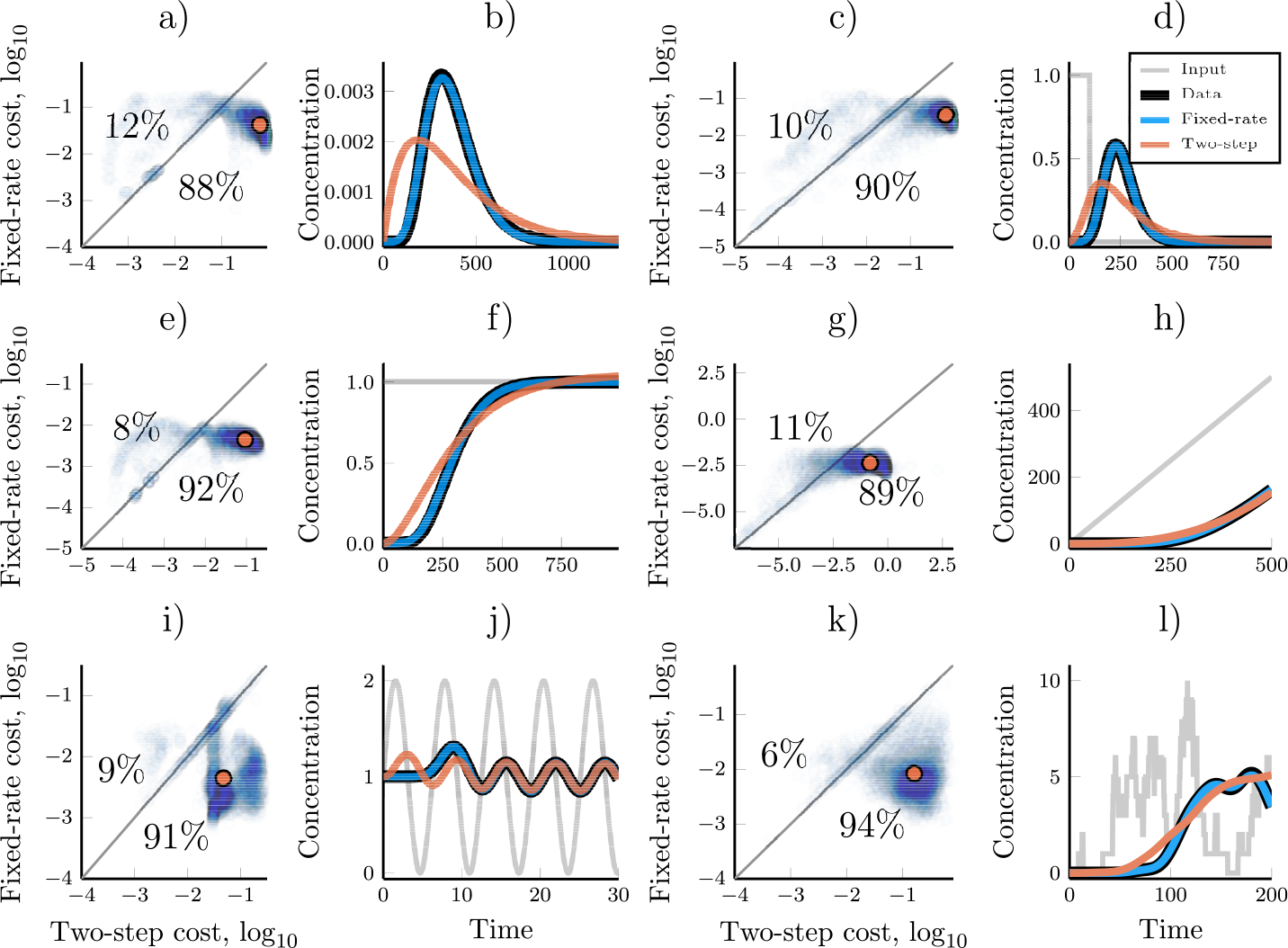
Comparing the ability of the fixed-rate and the two-step model to reproduce the dynamics of linear pathways which respond to different inputs. Each pair of figures (a,b; c,d; …; k,l) demonstrates the performance of the two models for the different model inputs: no input, piecewise constant, ramp, step, wave and noisy auto-activator, respectively (see Table 1). Figures a, c, e, g, i and k compares the optimised cost values (lower is better) for the fixed-rate model and the two-step model for each synthetic data set (5000 per input). Every data set is represented by a low-opacity dot; colour saturation thus indicate the density of similar values. Percentages indicate how often one model had a worse cost than the other. The orange dot shows the geometric median of the cost values for the different data sets. Figures b, d, f, h, j and l shows the model and data dynamics for the median data set of the corresponding input. The two-step model was chosen for the comparison since it has the same number of free parameters as the fixed-rate model.

**Table 1:**
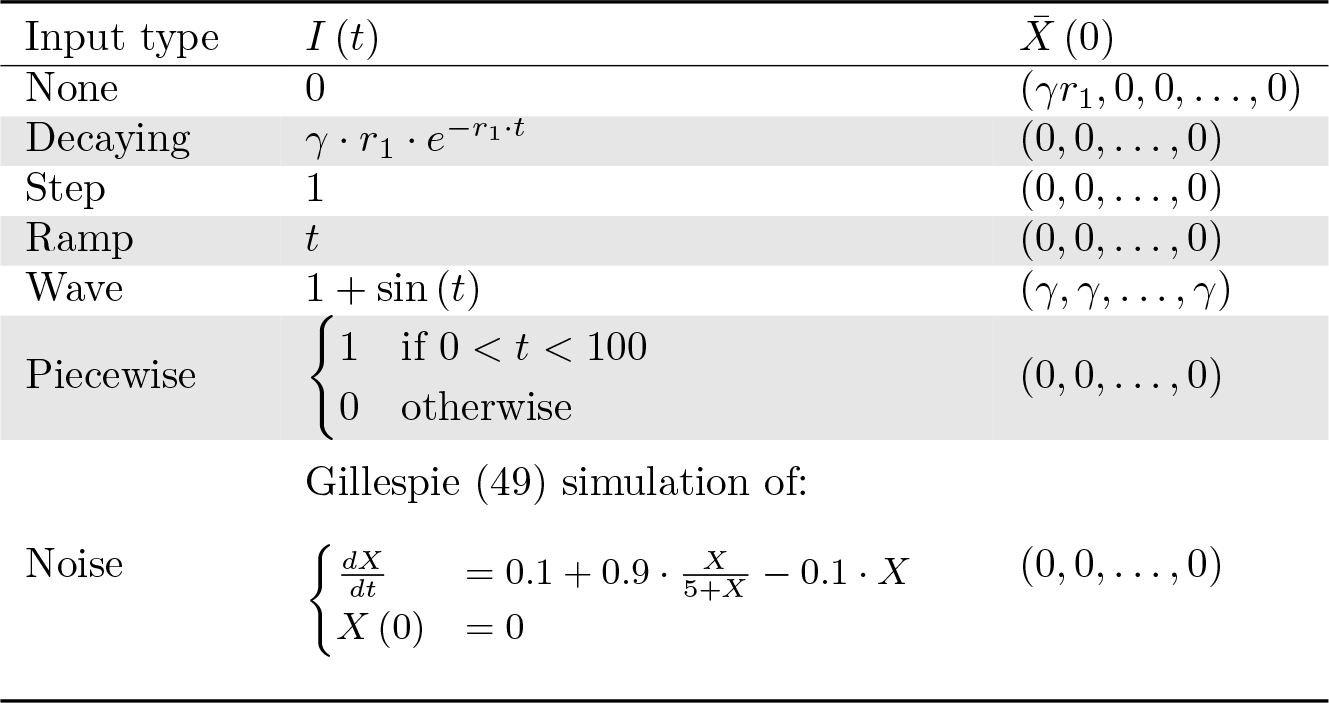
Model definition of linear pathways with different inputs. The models are all defined by Eqs. 2-3, with inputs, *I*(*t*), and initial conditions, *X̅* (0), as specified here.

A main limitation for the truncated models is that they perform badly in capturing the delay of a signal output. Rather than introducing multiple pathway steps in a model in order for it to capture a time-delay, such time-delays can be introduced explicitly. We next aimed to compare such an approach of modelling linear pathways to the use of the fixed-step and the fixed-rate formulations. In order to make a fair comparison, we defined a DDE with three free parameters. In this ‘fixed-delay’ model, the last pathway step, *X*_*n*_, responds to an input at a rate governed by *r*, with a fixed time delay *τ*. Similarly to the other models, a scaling parameter *γ* is also defined.

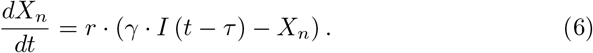

We optimised the fixed delay model (Eq. 6) towards the same data used for the previous models and compared their performance (Fig. 6). In most cases, this model performed better than the two-step truncated model but worse than the fixed-rate model. While this DDE model is better able to achieve a delayed response than the truncated model, it has issues with that response being too abrupt. The fixed-rate model, on the other hand, is able to achieve the correct time-delay while also accurately smoothing out the signalling over time.

**Figure 6:**
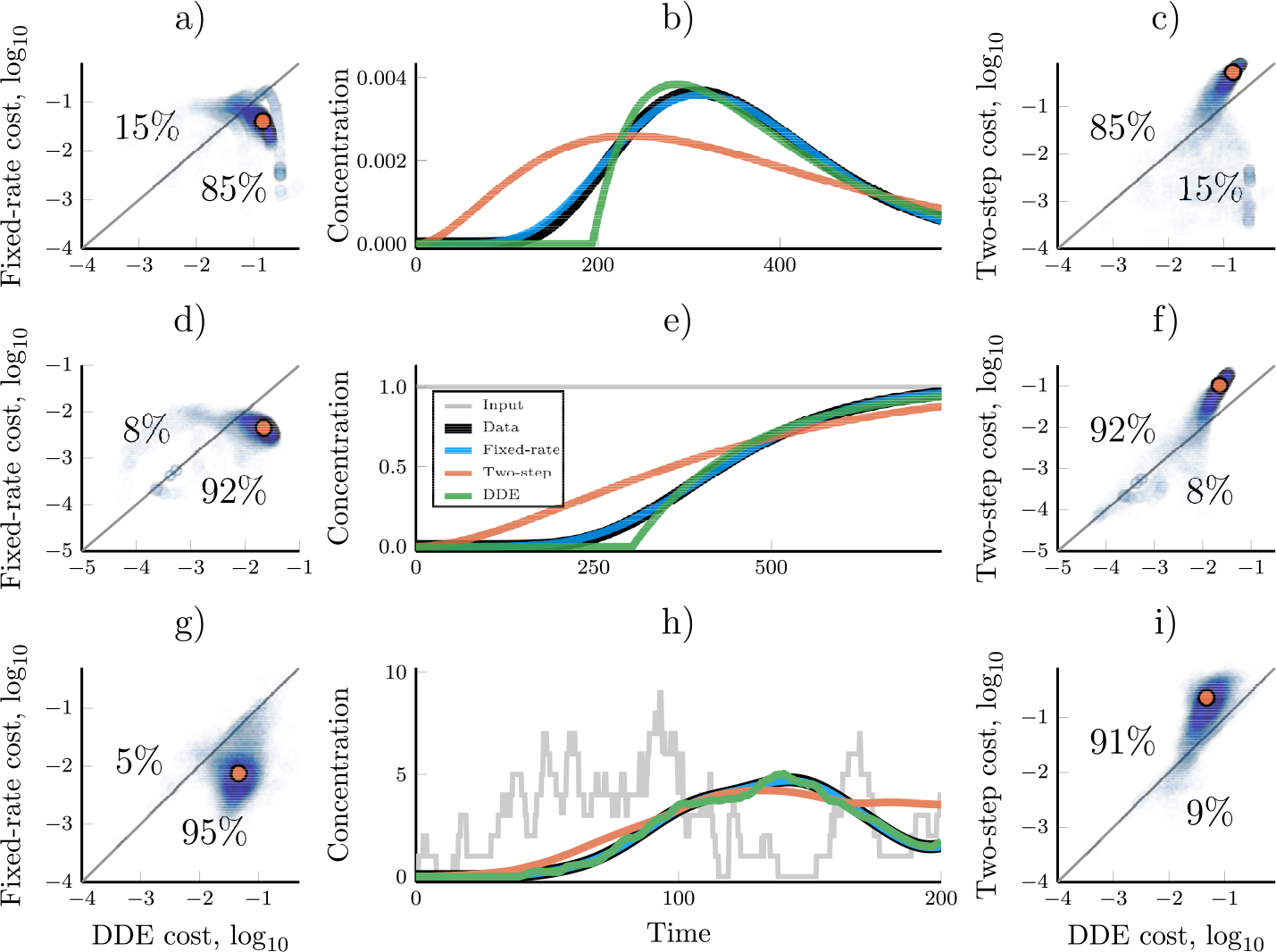
Modelling of linear pathways using a fixed delay DDE model (Eq. 6) compared to the two-step and the fixed-rate models. a) A cost value comparison of the fixed-rate model and the DDE model for every data set generated with a decaying input (Tab. 1, 5, 000 data sets). Every synthetic data set is represented with a low opacity dot; color saturation thus indicate a high density of similar values. The orange dot highlights the geometric median and that median data set is used as an example in figure b). b) An example time trajectory where the fixed-rate, two-step and DDE models have all been optimised to reproduce a synthetic data set. c) A cost value comparison, similar to a), between the two-step model and the DDE model. The orange dot shows the cost of the trajectory displayed in b). d-f and g-i) Repeats of a-c) but using a step input and a noisy input, respectively (Tab. 1). The synthetic data was generated with pathway lengths, *n*_*data*_, uniformly distributed between 1 and 50.

### Identifiability of biological parameters using the fixed-rate modelling approach

It is of interest to analyse how well a fixed-rate model approach performs when it comes to identifying the values of the parameters in the underlying biological pathway. When a pathway is using the same reaction rates for all pathway steps, the fixed-rate assumption is exact, and the parameter optimisation always finds the correct values for the underlying parameters. However, the model performs well even when representing a pathway with heterogeneous reaction rates, and in this case the connection between the model parameters and the biological system is less clear.

The parameter *γ* is a simple scaling parameter which was always set to 1 when we generated the synthetic data sets. Unlike for the fixed-step model (Eqs. 2-3), the optimised values of *γ* for the fixed-rate model were all very close to the true value (Fig. 4j). Both models are capable of ensuring that the signalling is properly scaled between the input and the output. However, since the fixed-step model is unable to correctly time its output, the optimisation scheme will sometimes lead to the use of the scaling parameter to mitigate the cost that this timing discrepancy creates. Since the fixed-rate model has much better control over timing, this never became an issue during our study, and the model always identified the correct value for the scaling parameter.

The parameter *n* represents the number of steps in the approximated pathway, but since we relaxed the demands that all rates are equal the connection between *n* and the number of steps in the biological (synthetic) data is not exact. This parameter value does not predict precisely the length of the linear pathway, but it does indicate a lower bound of the real pathway length (Fig. 4k). While the predicted *n* seems to scale linearly with the length of the pathway, the proportionality constant is dependent on the distribution from which the reaction rates of the linear pathway are drawn. If these rates are (close to) homogeneous, *n* will closely correspond to the number of steps in the system, while if the rates are more heterogeneous, the *n* will underestimate the length of the underlying pathway (Fig. 2f).

The parameter *r* of the fixed-rate model is related to the rates at which information is being passed along the linear pathway. The optimised value of this parameter cannot represent all rates in the pathway if these are heterogeneous. Instead, the *r* parameter seems to approximately identify the slowest part of the pathway since it is generally only slightly larger than the slowest reaction rate in the linear pathway (Fig. 4).

Altogether, there is not a perfect identifiability of the underlying pathway parameters when the fixed-rate model is optimised. Still, when analysing the resulting parameter values, strong indications of the parameter values and pathway structure of the original model can be found.

### Analytical solutions to the fixed-rate model extends its usefulness and dynamical range

For some simple inputs, the fixed-rate assumption allows for a concise and numerically efficient analytical solution to the problem (27). The analytical solution to the fixed-rate model with *I* (*t*) = 0 and *X̅* (0) = (*γ* · *r*, 0, 0, …, 0) yields a scaled version of the probability density function (PDF) of the gamma distribution (Methods, 27, 47), given by

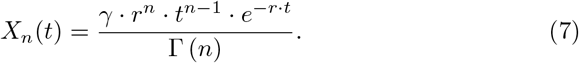

Because of its close connection to the gamma distribution, we will refer to this as the ‘gamma model’. In this model, Γ is the gamma function (47) and, just like in the fixed-rate model, *γ* is a scaling factor, *n* relates to the pathway length and *r* to the response rates of the steps along the pathway. For Eq. 7 to be an analytical solution to the fixed-rate model, *n* has to be an integer. However, while the fixed-rate model cannot account for partial steps, the gamma model can.

Relaxing the demand that *n* is an integer increases the freedom of the model and thus possibly its ability to fit data compared to the fixed-rate model. To test the analytical model for pathways with arbitrary reaction rates, we optimised the fixed-rate model and the gamma model towards the same synthetic data sets (Methods). The gamma model, with *n* ∈ [1, ∞), improved performance compared to the fixed-rate model in nearly all cases (Fig. 7, cf. Fig. S5, 27). In most cases, the two models were approximately equivalent but the performance difference became apparent in some cases, especially when *n*_*data*_ was small.

**Figure 7:**
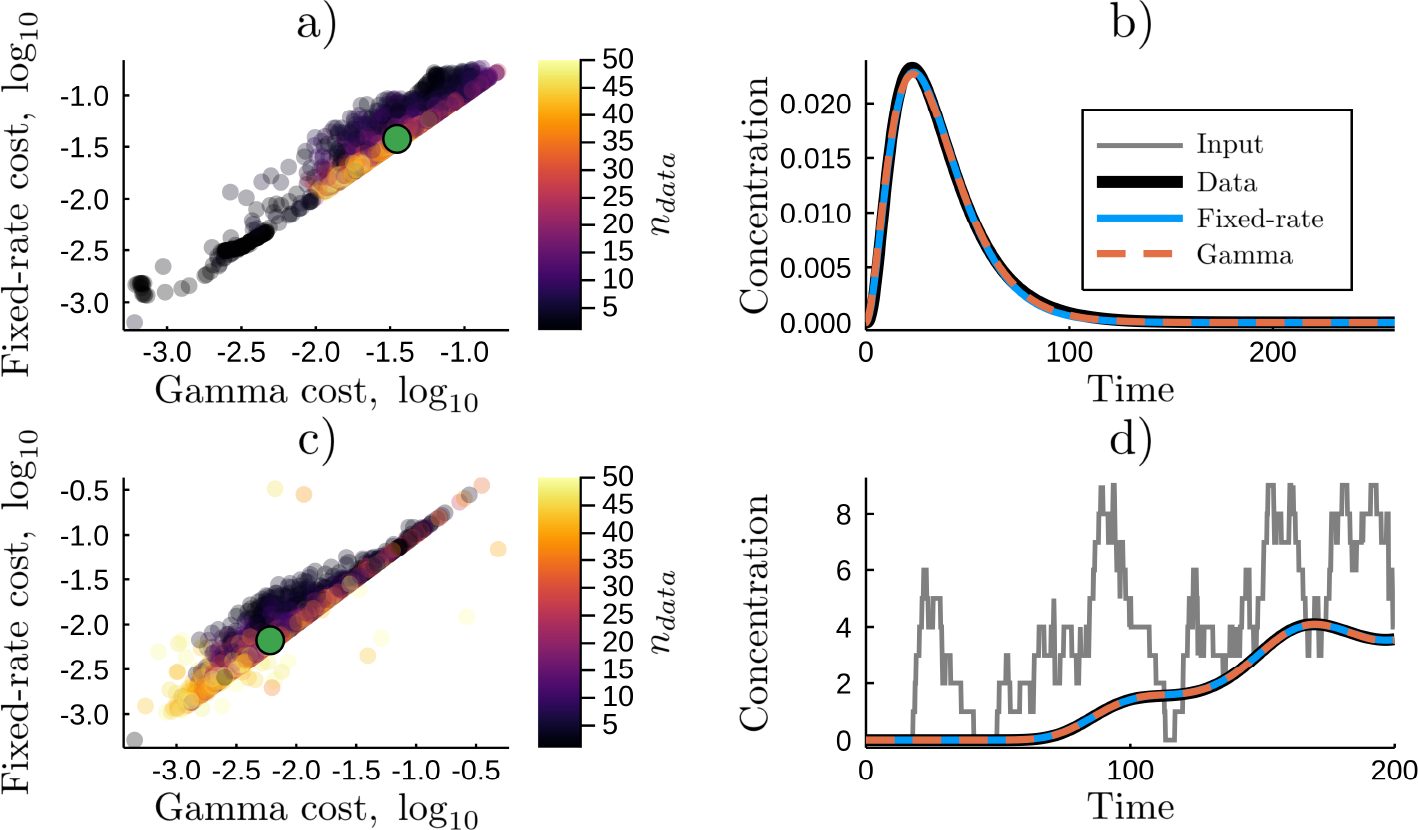
Allowing a real-valued pathway length parameter, *n*, increases the gamma model’s ability to recapitulate dynamics from arbitrary linear pathways. The optimised cost values for the fixed-rate and the gamma models when they are both optimised towards the same data sets in which the pathway has no input (Tab. 1, 5000 data sets). The colour of the dots indicate the length of the linear pathway which was used to generate the synthetic data set. b) Demonstrating the model performances for the median data set, as denoted by a green dot in (a). c, d) similar to a and b but using the generalised gamma model (Eq. 8) and a noisy input (Tab. 1).

Many of the gamma model’s characteristics can be inferred directly from its parameter values without needing to simulate the model. Since the gamma model is closely connected to the gamma distribution, some of the characteristics and statistics of the distribution are directly applicable to the model. The mean of the gamma distribution identifies the time, 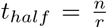, at which half the signalling will have occurred for *X*_*n*_(*t*), the variance is given by 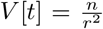 and gives a measure of how the signalling peak is spread out over time. This directly identifies that if *n*_*model*_ < *n*_*data*_ the model peak must be more spread out if the time of the peak is to match that of the data. The mode gives the time at which the distribution/model reaches its peak value and is given by 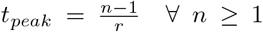. If *t*_*peak*_ is a main target of the model fitting, this relation can be used to reduce the size of the parameter space search.

The gamma model can be extended to allow for any integrable input. The convolution between the gamma model and the input results in a gamma distributed delay model (Methods).

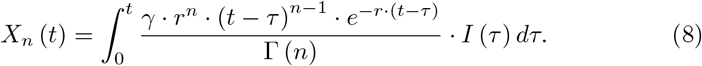

This generalisation of the gamma model allows for (nearly) arbitrary inputs and thus greatly enhances the scope of the model. This, for example, enables its use with dynamical inputs or within feedback loops inside of DDEs. The downside to this formulation is the need to evaluate an integral which is computationally costly. The model is still restricted in the possible initial concentration of the pathway where it is applicable since its derivation assumes an all zero initial concentration along the pathway. However, this restriction can often be circumvented. By setting the start-time of the integral to be before the start time of the actual simulation, it is possible to equilibrate the solution to some arbitrary input before the simulation starts (Methods).

## Discussion

Biology is full of multi-step pathways where each individual step is (at least approximately) linearly dependent on the previous step. Transcription, translation, kinase cascades, sequential phosphorylation and signal transduction are all examples of processes where this can apply. The full inclusion of such pathways is seldom advisable when modelling biological systems since the added benefit in dynamical range is outweighed by the disadvantage of an increased model complexity. It is, therefore, common for such pathways to be simplified. However, the manner in which such pathways are simplified is not always particularly effective.

We demonstrate how the coarse-graining of a long linear pathway to a short one (pathway truncation) often lead to a detectably incorrect temporal relationship between the input and the output signal. This discrepancy is important to understand not only because it can cause a model to quantitatively misrepresent time-course data but also because signal timing can qualitatively alter dynamical behaviour. Negative feedback loops, for example, can change from having a stabilising effect to generating oscillations when the feedback is delayed (e.g. 1, 11). It is, therefore, notable that when this simplification has adverse effects we could identify a detectable signature in the form of homogeneity of the optimal response rate parameters. This can be used as a model diagnostic and could prove especially helpful for complex models where the source of model/data mismatch is not always apparent from their design or output.

Next we asked whether there might be a way to remedy the shortcoming of a truncated model without increasing the number of model parameters. Our proposal is to assume that the signalling is being passed along the pathway at a constant rate while the number of steps in not fixed. This assumption allows for a three-parameter approximation of arbitrary linear pathways where the pathway length is a tunable parameter. We showed that this ‘fixed-rate’ assumption clearly outperformed both truncated pathway models and a DDE model with an explicit time-delay parameter even if the original pathway had highly heterogeneous rates for individual pathway steps. It also naturally leads to a gamma distributed delay model where analytical solutions can be tractable (see e.g. Beguerisse-Díaz et al. 27).

For a model to be useful, it is important to retain information about the underlying biological system even after the model assumptions have simplified reality. Much of the connection between the model parameters and real biological processes is retained when the fixed-rate assumption is used for simplification of a linear pathway. The pathway length parameter of the fixed-rate model, *n*, provides a rough lower bound for the length of the real pathway. Also, the value of its response rate parameter, *r*, is strongly correlated with the response rate of the slowest step in the underlying system. This should be contrasted with the truncated pathway model which still uses individual response rate parameters for individual pathway steps. Intuitively, this may seem more closely related to reality. However, when such a model is optimised to perform the action of a longer pathway, these individual reaction rates become highly decoupled with the response rates of any real pathway step. Rather than being tuned to represent real pathway steps, they are tuned to delay the response peak while not sacrificing too much of the response sharpness. So the fixed-rate assumption not only performs better than the alternatives, its individual components also retains a closer connection with reality.

In the future it will be interesting to understand if the performance increase in predicting a pathway’s input-output dynamics will hold also when a linear pathway is part of a larger dynamical system, including multiple interactions and feedbacks. Recently, Tokuda et al (43) showed that introducing distributed delays (fixed-rate assumption) between nodes in circadian clock models allowed for a parameters reduction. Given that they did not allow for fully dynamic changes of pathway lengths (*n*), it would be interesting to see if this can improve the model predictability. We limited the scope of this paper to that of deterministic systems, only introducing stochasticity in the input to the pathway, but it would also be interesting to see how the reasoning applies in a fully stochastic description. Similarly, relaxing the linearity constraint in the underlying models would provide an additional interesting challenge for the suggested fixed-rate models that can be analysed.

A mathematical model is defined by its underlying assumptions and can be seen as merely acting as a logical device to deduce consequences of those assumptions (48). A main part of model development is thus to find a set of assumptions which accurately and concisely captures the nature of the dynamical system under study. However, finding a good balance between detail and simplicity is often non-trivial and requires some degree of craftsmanship, and the better we understand the consequences of specific assumptions, the better we become at striking this balance. In this context, we systematically investigate the consequences of simplifying linear pathways by truncating the number of steps in a pathway. We clearly demonstrate problems such simplifications may cause. More importantly, we present how to detect these issues when they occur and we provide an alternative approximation to remedy them. We hope this will supply a foundation for well-informed decisions regarding when and how to simplify the ubiquitous linear pathway.

## Methods

### Model derivation

If we model an *n*–step linear pathway that follows Eq. 1 except that it receives some input signal to the first step, *X*_1_, the dynamical equations can be written as

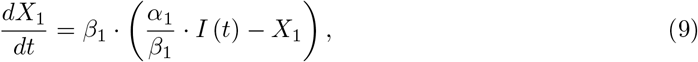

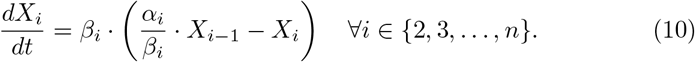

where *α*_*i*_ is a production/activation rate, *β*_*i*_ a degradation/deactivation rate and *I* (*t*) is some upstream input. Assuming that the input is integrable, the Laplace transform, 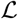, of this system is (27, 47)

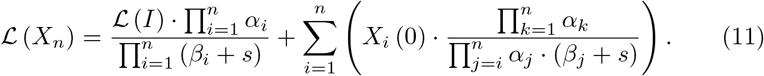

Here, the first term describes the systems dynamics from rest, *X̅* = (0, 0, …, 0), and the second term compensates for the initial concentrations of the pathway (27). This tells us that if the pathways starts from rest then all the production terms, *α*, are linearly dependent. Utilising this, we define *α*_1_ = *β*_1_ · *γ* and *α*_*i*_ = *β*_*i*_ ∀_*i*_ ∈ {2, 3, … , *n*} so that we can use the single parameter, *γ*, to scale the output. Changing the ndividual *α*_*i*_ values would have changed the effect that non-zero initial concentrations along the pathway would have. However, this effect is itself linearly dependent on the actual initial concentrations *X̅* (0). A transformation of *X̅* (0) can therefore compensate for our parameter reduction so that no dynamical range is lost at all for the model’s input-output relationship. Since this simplification means that any change in the degradation parameter, *β*, no longer scales the output, we rename it *r* ≡ *β* and call it a response rate parameter to better reflect its action. The result is the linear pathway model we have been using in the paper, Eqs. 2-3.

The analysis of the models were made using different sets of inputs and initial concentrations (Table 1). With these definitions, a model of pathway length *n* with the ‘Decaying’ input is identical to models of pathway length *n*+1 with the ‘None’ input. The noisy input was generated using a Gillespie simulation of a single, auto-activating, variable (49). The noisy time-course that this generated was used as an input both during the generation of synthetic data and during the subsequent model simulations.

### Analytical solution of the fixed-rate model with no input

It is possible to find analytical solutions to Eqs. 4-5 for some simple inputs to the linear pathway. Here, we demonstrate this for the case of no input (Tab. 1) but the same has been done for more inputs in (27).

With *I* (*t*) = 0 and *X̅* (0) = (*γ* · *r,* 0, 0, …, 0), the fixed-rate model reads

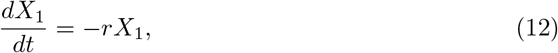

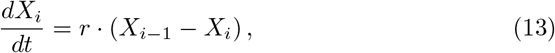

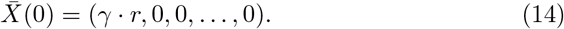

From here, we apply the Laplace transform, 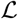, on both sides of Eqs 12-13, utilising the fact that the Laplace transform of a function derivative, *f*′, follows 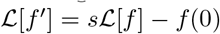. This leads to

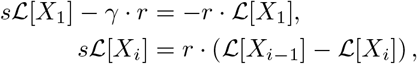

which can be rearranged to get

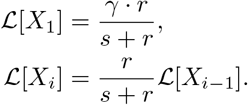

This can be recursed to solve for *n* steps leading to

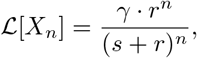

and the inverse transform of this yields

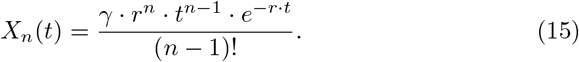

By expanding the domain of *n* to that of the real numbers, we get the gamma model (Eq. 7).

### Derivation of the gamma distributed delay from the fixed rate assumption

Laplace transforms and the transfer function provides a way of finding an analytical solution to the fixed-rate model with arbitrary inputs (47, 50).

The laplace transform of the fixed-rate model ODEs (Eqs 4-5), with initial condition *X*_*i*_ (0) = 0 ∀*i*, is given by

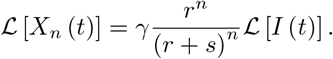

From this, we can easily get the transfer function in the complex domain

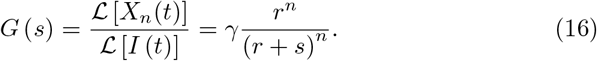

The transfer function in the time domain, *g* (*t*), is given by

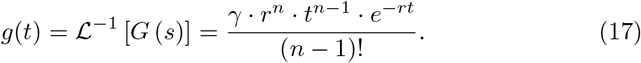

Here, we see that the transfer function in the time domain (also called the weighting function) is essentially the gamma model. This transfer function can be used to get an analytical solution to *X*_*n*_ (*t*) for almost any input (the input needs to *have* a Laplace transform, even if we never have to calculate it). From Eq. 16, we have that

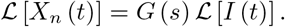

This equation is still in the complex domain but we can use the connection between multiplication in the complex domain to convolution in the time domain to get the desired function

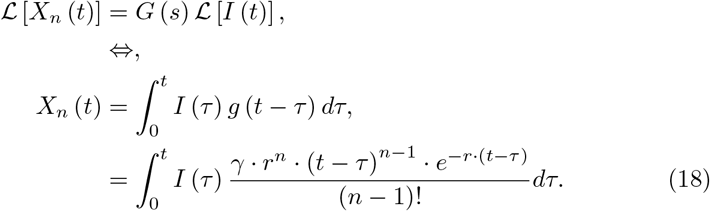

In order to improve the model’s ability to fit data, we can replace (*n* − 1)! with Γ (*n*). This makes no difference for integer *n* but it expands the possible domain of *n* to all real numbers greater than or equal to 1. After this, we end up with the gamma distributed delay, scaled with *γ*, of eq. 8.

### Model simulation

Differential equations were solved using an algorithm with stiffness detection that toggled between Tsitouras5 for non-stiff regions and Rosenbrock23 for stiff regions (51–53). The integral of the gamma distributed delay model (Eq. 8) was evaluated using Gauss-Kronrod quadrature.

For the wave input we needed an initial concentration of *X*_*i*_(0) = *γ* ∀*i*. For the gamma distributed delay model, this was achieved by using the input function

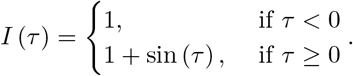

and by starting the integration at a negative *τ*. The actual value used was based on the mean and the standard deviation of the gamma distributions PDF

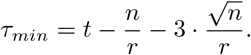

### Data generation

Synthetic data sets were generated using the fixed-step model (Eqs 2-3). The procedure was to first draw an integer between 1 and 50 to be used as the number of pathway steps, *n*_*data*_. A response rate, *r*_*i*_ was randomly drawn for each of the *n*_*data*_ steps in the pathway. For most input types, the parameters *r*_*i*_, were generated by transforming the uniformly random variable *Y*_*i*_ ~ *U* (−2, 1) according to 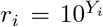. However, since any too slow reaction rates will filter out the dynamics of the noise and the wave input, the reaction rates for those models were drawn from 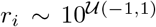 and 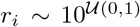, respectively. The value of the scaling parameter, *γ*, was in all cases set to 1. The model was then run and the resulting trajectory of the last pathway step, *X*_*n*_(*t*), was stored for use as the synthetic data set (5000 times for each input).

### Fitness definition

In order to automatically evaluate the fitness of a model, an ‘integral cost’ function was defined. The idea of this cost function is to measure the mismatch in the area under the curve for the model and the data (Fig. 8). This is similar to using the *ℓ*1 norm, but it has a few advantages: It does not require the differential equation solver to stop at the time-points where the data was sampled and variations in sample density does not bias the cost value. We defined this cost value as

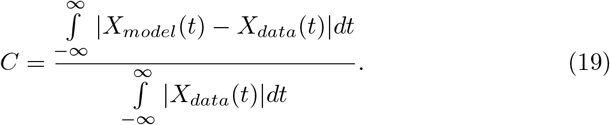

**Figure 8:**
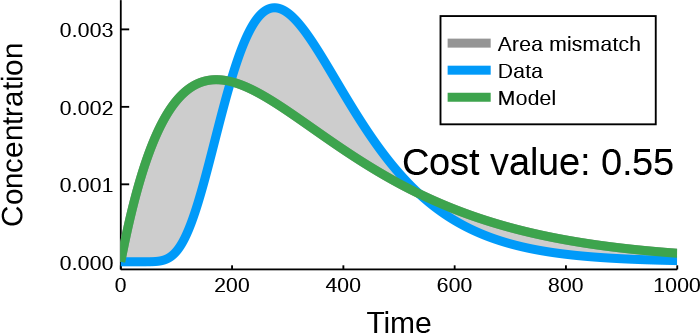
Cost value definition. The cost value measures the normalised area mismatch between the model output and that of the data. The area under the data curve is used for the normalisation.

Numerically, *C* was calculated using Riemann sums and interpolations of both the data and the model solution. The evaluations started at *t* = 0 and for the ‘None’, ‘Step’, and ‘Piecewise’ input types they continued until the model derivatives were close to zero (absolute and relatives tolerance 10^−8^ and 10^−6^, respectively) or until *t* = 5000, whichever came first. The ‘Piecewise’ input simulations were additionally prohibited from stopping before the end of the piecewise constant input. Since the ‘Ramp’, ‘Wave’ and ‘Noise’ inputs do not allow for equilibration, we set fixed stopping times of *t* = 500, 30 and 200, respectively.

Much of the analysis was also repeated using the more commonly used normalised least square cost function. While the results are slightly different, they did not change any of the conclusions in this work.

### Model optimisation

The model parameters were all optimised to reproduce the time-trajectory of each synthetic data set. The optimisation target was to minimise the cost value, *C*, described above. For the actual optimisation, we used an adaptive differential evolution algorithm following the /rand/1/bin/ scheme, with a radius limited sampler that took 2,000 steps per free parameter of the model (54, 55). The search space for the parameters of the different models were chosen to allow for the time delays that the synthetic data could generate (Tab. 2). The sampling space was linear for *n* and logarithmic for the other parameters.

**Table 2:** Parameter search space for the optimisation of the different models.

| model | $\gamma$ | $r$ | $n$ | $\tau$ |
| --- | --- | --- | --- | --- |
| fixed step | $[10^{-3}, 10^3]$ | $[10^{-3}, 10^3]$ | N/A | N/A |
| fixed rate | $[10^{-1}, 10^1]$ | $[10^{-2}, 10^2]$ | $\{1..30\}$ | N/A |
| gamma | $[10^{-4}, 10^2]$ | $[10^{-3}, 10^3]$ | $[1, 31]$ | N/A |
| DDE | $[10^{-4}, 10^2]$ | $[10^{-4}, 10^1]$ | N/A | $[10^{-2}, 10^4]$ |

### Software availability

All the computational results were generated using the Julia programming language (56). Differential equations were solved using DifferentialEquations.jl, Gauss Kronrod quadrature was done with QuadGK.jl and optimisation was done with BlackBoxOptim.jl (53, 55). The source code developed for this project is openly available under the MIT licence at the Sainsbury Laboratory GitLab repository https://gitlab.com/slcu/teamHJ/publications/Korsbo_et_al_2019. This repository also includes documentation and a tutorial aimed at making reproduction and reuse easy.

The software for the evaluation of the integral cost function can also be accessed independently at https://gitlab.com/slcu/teamHJ/niklas/CostFunctions.jl.

## Acknowledgements

We would like to thank Torkel Loman and members of the Jönsson group for fruitful discussions and Torkel Loman, Ross Carter and James Locke for feedback on the manuscript.

This work was supported by the Gatsby Charitable Foundation, grant GAT3395-PR4.

