## Supplemental Information for "It’s about time: Analysing an alternative approach for reductionist modelling of linear pathways in systems biology"

### Supplementary Information

Niklas Korsbo and Henrik Jönsson

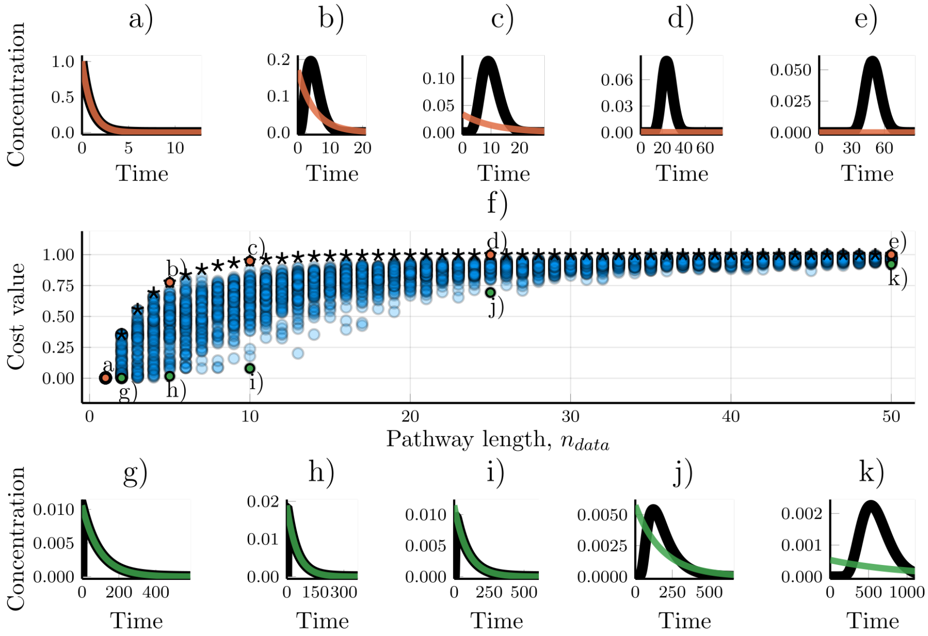


Figure S1: Fitting one-step models to data from linear pathways of different length. a-e) The worst model/data fits for a given length, $n_{data}$, of the model that generated the data. Coloured lines show simulations of the fitted model while black lines show the synthetic data. (f) The cost value for 5000 parameter sets, each optimised towards a different set of synthetic data. Circles show the cost values resulting from data wherein all the steps in the data-generating linear pathway were randomly drawn, $r_i \sim 10^{U(-2,1)} \quad \forall\ i$ . Stars are the cost values from data wherein the pathway has homogeneous reaction rates, $r_i = 1 \quad \forall\ i$. The x-axis shows the number of steps in the model which were used to generate the data. g-k) Examples of the best model/data fits for different data pathway lengths, $n_{data}$.

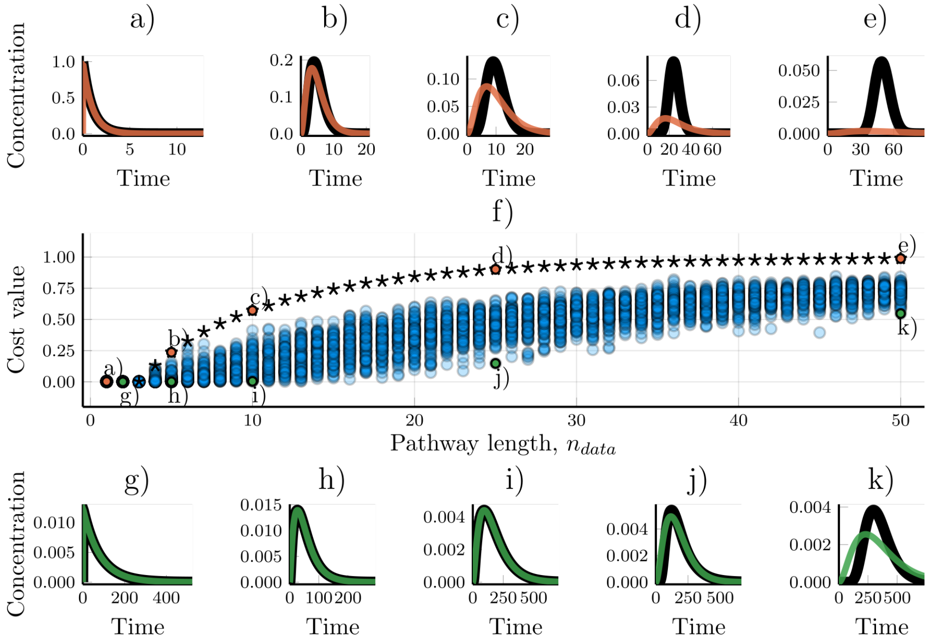


Figure S2: Fitting three-step models to data from linear pathways of different length. For a longer description, see the caption of Fig S1.

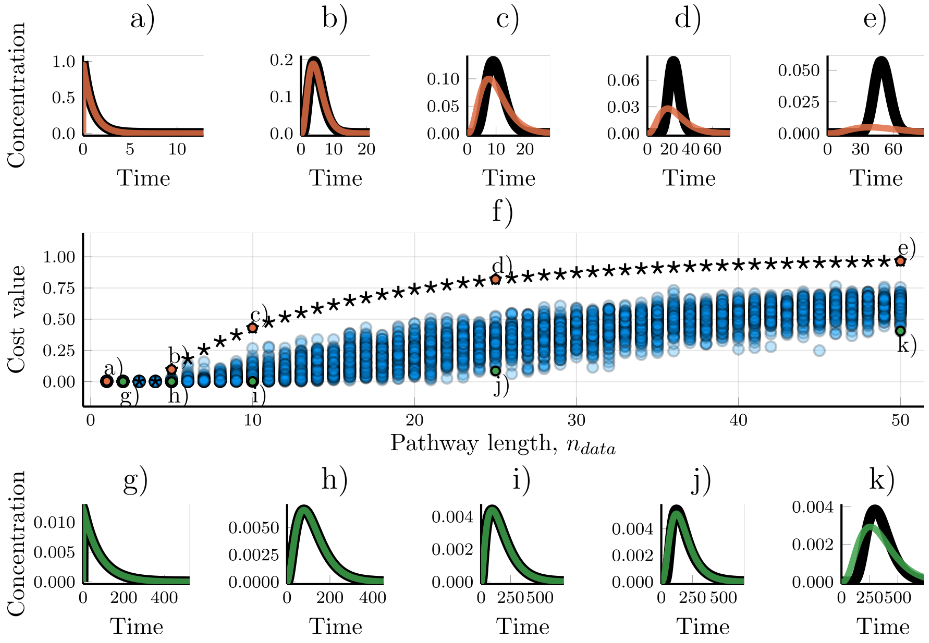


Figure S3: Fitting four-step models to data from linear pathways of different length. For a longer description, see the caption of Fig. S1.

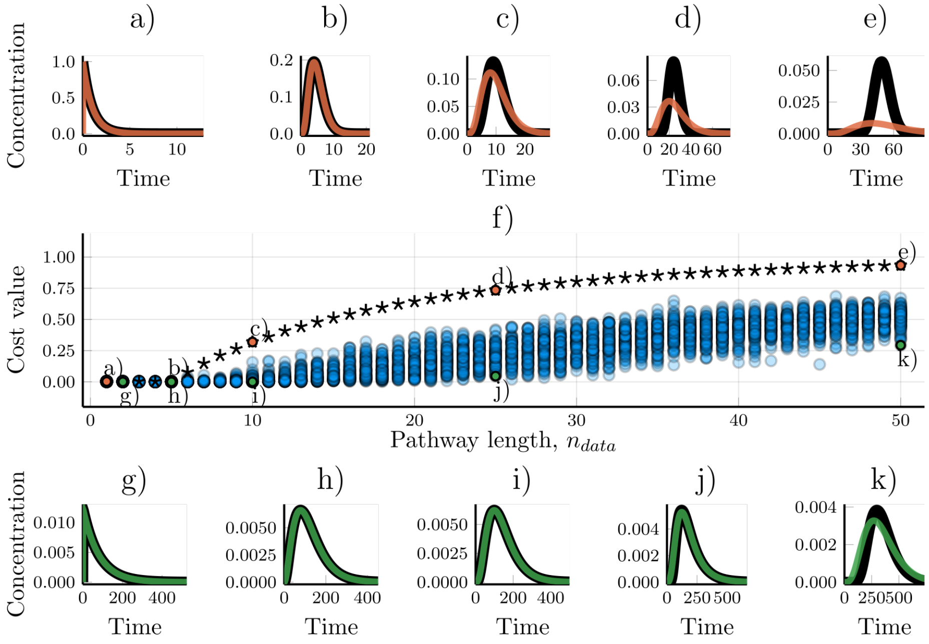


Figure S4: Fitting five-step models to data from linear pathways of different length. For a longer description, see the caption of Fig. S1.

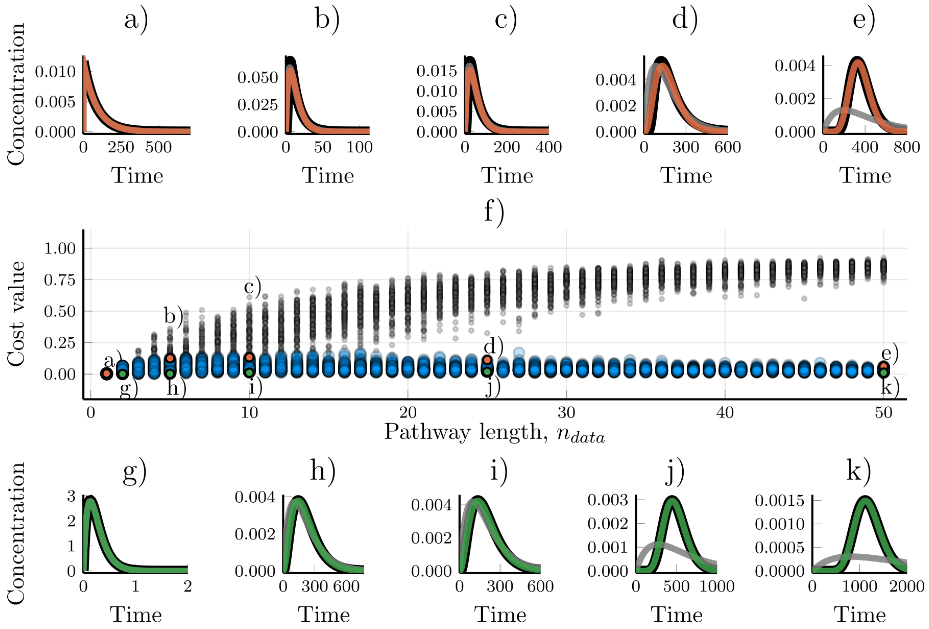


Figure S5: Fitting gamma models to data from linear pathways of different length. The gamma model (Eq. 7) with $n_{model} \in \mathbb{R}$ efficiently recapitulates linear pathway dynamics. a-e) Examples of the worst model-data fits for different lengths of the data-generating pathway, $n_{data}$. Black lines show the synthetic data, orange lines show the results of the gamma model, and grey lines show the results of a two-step model (Eqs. 2-3). f) The ability of the gamma model to fit the data changes with the length of the underlying pathway, $n_{data}$. Blue dots show the cost value of the gamma model. The small, grey, dots show the corresponding cost values for a two-step model. g-k) Examples of the best model-data fits.

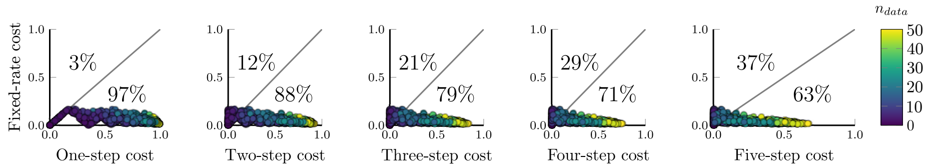


Figure S6: A comparison of the cost values when using either the fixed-rate model (Eqs. 4-5) or a fixed-step model (Eqs. 2-3) to fit the same data. Each circle represents a single synthetic data set. 20 synthetic data sets were generated for each $n_{data} \in \{1, \ldots, 50\}$ using the 'None' input (Tab. 1). The circles position indicates the optimised cost for the respective models and its color indicates the $n_{data}$ value of the data-generating pathway. Percentages indicate how many of the data sets had a higher (worse) cost value for the respective models.
